# Phenotypic and genetic heterogeneity of *Acinetobacter baumannii* in the course of a chronic infection

**DOI:** 10.1101/2024.05.22.595333

**Authors:** Léa Bednarczuk, Alexandre Chassard, Julie Plantade, Xavier Charpentier, Maria-Halima Laaberki

## Abstract

*Acinetobacter baumannii* is a nosocomial pathogen associated with various infections, including urinary tract infections (UTIs). In the course of an infection, *A. baumannii* is known to rapidly become resistant to antibiotic therapy, but much less is known about possible adaptation without antibiotic pressure. Through a retrospective study, we investigated within-host genetic diversity during a subclinical five-year UTI in an animal-patient after withdrawal of colistin treatment. We conducted whole-genome sequencing and phenotypic assays on seventeen clonally related isolates from the Sequence Type 25 lineage. Phylogenomic analysis revealed their proximity with animal and human strains from the same country suggesting zoonotic transmission (France). In this case study, the clonally related strains presented variations in genome sizes and nucleotide sequences. Over the course of the infection, *A. baumannii* underwent genome reduction through insertion sequence (IS) recombination, phage excision, or plasmid curing. Alongside this global genome reduction, we observed an expansion of IS*17*, initially located on the endogenous large plasmid. Genetic variations were mainly located in biofilm formation and metabolism genes. We observed repeated variation affecting three biofilm genes and two adhesion operons associated with weak biofilm-forming capacity. Conversely, only two metabolic genes were recurrently affected and phenotypic assays indicated a rather stable metabolism profile between the isolates suggesting minor adaptations to its host. Lastly, an overall decreased antibiotic resistance – expected in the absence of antibiotic treatment - contrasted with a conserved colistin resistance due to a *pmrB* mutation among the isolates.

**Impact statement:** This study brings a new insight on the genome evolution of the opportunistic pathogen *Acinetobacter baumannii* during a five-year urinary tract infection in the absence of antibiotic treatment. It relies on genomic and phenotypic analyses of multiple isolates from the same animal-patient highlighting the necessity of studying bacterial diversity as opposed to solely on single-isolate approaches. Over the course of the infection, we observed a reduction in genome size, resulting however in a limited loss of metabolism flexibility, but we observed recurrent loss in adhesion capacities and antibiotic resistance. However, this study stands out by the unexpected conserved colistin resistance of the study isolates in the absence of treatment.

**Data summary:** The newly sequenced genomes are listed in Table S1, these genomes have been submitted to the NCBI and their BioProject number is PRJNA1100485. The publicly available genomes of bacteria from human and animal origin used in this study are listed in Table S2 All the supplementary tables are available in the Supplementary Material File.

## Introduction

*Acinetobacter baumannii* is a non-fermentative Gram-negative pathogenic bacterium responsible for nosocomial infections in humans and animals, particularly in intensive care units (1, 2). *A. baumannii* represents a major threat to public health due to the rapid emergence of multidrug-resistant isolates increasingly difficult to eradicate (3). Indeed, few antibiotics remain effective against this pathogen, such as carbapenems, and the carbapenem resistance of clinical isolates may require poorly tolerated antibiotics such as colistin (4). Emergence of carbapenem resistance in *A. baumannii* therefore limits treatment option and led World Health Organization (WHO) to classify carbapenem-resistant strains of *A. baumannii* (CRAB) as a priority for research and development of new drugs (5). This pathogen is also described in animal infections especially in pets with strains belonging to clones common with human medicine with concerning resistance to human restricted antibiotics such as carbapenems (1, 6).

This bacterium has no particular tropism, causing mainly pulmonary infections but also capable of skin, blood, bones, urinary tract infections (7). The existence of extra-hospital reservoirs has long been disputed, but recent studies demonstrate that *A. baumannii* is an ubiquitous bacterium found both in soils and carried by healthy individuals - humans and animals (8–11). This bacterium is therefore capable of thriving in highly contrasted environments, demonstrating a great capacity for adaptation; antibiotic resistance would be just one facet of its versatility applied to the healthcare context.

Several within-host studies of *A. baumannii* infection describe antibiotic resistance emergence in response to treatment due to non-synonymous single nucleotide polymorphism (SNP) or horizontal gene transfer (12, 13). However, little is known about host adaptation, because brief timeframes of infection limiting observation of bacterial evolution within a host. Additionally, these investigations focused on human infections, typically involving extended antibiotic treatments, making it difficult to decipher bacteria’s adaptation to the antibiotic from host adaptation. They also involve mostly single-isolate approaches hindering the understanding of within-host diversity. Consequently, there remains a limited understanding of the fate of antibiotic resistance genes and more broadly of gene evolution post-antibiotic exposure and during host colonization. Seminal within-host microevolution studies on *A. baumannii* observed the accumulation of variations in genes involved surface structures such as motility, adhesion, biofilm formation or cell wall and capsule biogenesis, suggesting a consistent selective pressure during antibiotic-treated infections but phenotypic validation are usually missing (14–16). Phenotypically, antibiotic exposure prompts bacteria to adopt a biofilm-like lifestyle, enhancing their resistance (17). These biofilms pose challenges for infection treatment and promote recurrence, notably in urinary tract infections (UTI) (18). Despite the over-representation of pulmonary and bloodstream infections in *A. baumannii* studies, the urinary tract is a major source of *A. baumannii* isolates in human (19). Tissue tropism may result from mobile genetic elements (MGE) regulation as exemplified by plasmid-driven regulation of chromosomal genes modifying strain adhesion pattern (6, 20, 21). Therefore, *A. baumannii* within-host microevolution may involve crosstalk of MGE and core genome determinant.

In the following case study, a pet infection emerged from a subcutaneous ureteral bypass placement which was particularly described as a risk factor for development of *A. baumannii* UTI in cats (22). We used whole-genome sequencing and phenotype assays on 17 *A. baumannii* strains isolated from the same cat with UTI over a five-year period without antibiotic treatment. We sought to investigate the fate of antibiotic resistance and explore evolutionary strategies upon infection.

## Materials and methods

### Strains

*A. baumannii* strains identified in the Laboratoire Vétérinaire Départemental du Rhône, France (LVD69) were stored in an archive since 2014. Nine strains originated from distinct urine samples collected between 2014 and 2019 and referred to as the longitudinal strains, while all others were derived from a single urine sample collected later in October 2019, referred to as the synchronic isolates. Samples were obtained not for the purpose of this study or research, but rather for diagnostic purposes. The longitudinal strains were initially isolated on a blood agar plate and the synchronic strains were isolated using three different media: LB (Lysogeny Broth, Sigma) agar, Mueller-Hinton agar (Sigma) and CHROMagar *Acinetobacter* (CHROMagar). All the strains used in this study are listed in Supplemental **Table S1**.

### Metabolic assays

Metabolic assays were conducted using Biolog Gen III Microplates (Hayward, CA) following the manufacturer’s instructions. In brief, the strains were streaked on LB-agar and incubated over night at 30°C. For each strain, a single colony was resuspended in inoculating fluid A (IFA, Biolog) using a cotton-tipped Inoculatorz swab (Biolog) to reach a cell density of 90-98%. Each well of the Gen III MicroPlate, containing either a specific carbon source or a chemical sensitivity testing agent, were filled with 100 μL of the bacterial suspensions and incubated at 33°C for 48 hours in the OmniLog incubator/reader measuring the reduction of tetrazolium redox dye (forming a purple color) due to bacterial growth.

### Genome sequencing and annotation

Bacteria were grown in liquid LB medium and DNA was purified using Wizard genomic DNA kit (Promega). DNA quantifications were performed using Qubit™ dsDNA HS Assay Kit (Thermo Fisher Scientific). For each isolate, Oxford Nanopore and Illumina DNA sequencings were both performed. Nanopore sequencing were performed using MinIon flow cells [FLO-MIN106] using the SQK-RBK004 barcoding kit. Illumina sequencing was performed on Illumina HiSeq 2000 with paired 150-base sequence reads (Novogene). Each read set was assembled individually using Unicycler (version 0.4.9, 41) and annotated using Prokka (version 1.14.5, 42) on the Galaxy pipeline using as reference AB5075-UW protein sequences (GeneBank accession number: CP008706.1). For analysis purposes, plasmid pF14-11 was annotated using pA297-3 sequence (GeneBank accession number: KU744946) except for the *tet*R genes that were named after pAB04 (Genebank accession number: CP012007, 19, 49–52). Putative prophages were predicted using PHASTER (27). Putative resistance genes were detected using Resfinder (version 4.5.0, 54) and insertion sequences using ISfinder (29). All genomes are available at NCBI under BioProject PRJNA1100485 with quality performed by the NCBI platform using CheckM (v1.2.2) with completeness between 98.46% and contamination ranging from 0.27% to 0.53%.uu

### Construction of bacterial strains

All the oligonucleotides used in this study for genetic modification are listed in supplementary **TABLE S3**. Gene disruptions were performed using a scarless genome editing strategy described previously (30). For plasmid curing of strain F14-11, its Δ*higA-higB::sacB-aac4* derivative was grown overnight in LB (2mL) at 37°C, the next day, 100µL were spread on M63 with 10% sucrose for 24h growth at 37°C then purified on the same medium. About 150 sucrose-resistant CFUs were then patched on LB agar, LB Tetracycline 30µg/mL, LB Apramycin 30µg/mL. Tetracycline and apramycin resistant clones were screened by PCR and plasmid extraction for plasmid loss.

### Phylogenetic analysis of the study strains

All chromosomes were aligned to the F14-11 chromosome, set as a reference, using the Gubbins script generate_ska_alignment.py, which creates an alignment using split k-mer analysis version 2 (SKA2, version 0.3.7, 58). The resulting alignment measured 4,061,760 bp with 337 variable positions. The strain 17021 was additionally included to root the tree. This strain is a suitable root as it lies outside the other study strains while still being phylogenetically close. Maximum likelihood tree reconstruction was performed with Gubbins (version 3.2.1, 57) relying on RAxML using the GTRGAMMA model (version 8.2.12, 56) for constructing the phylogeny in each iteration to mitigate the bias of horizontal sequence transfer mechanisms. Statistical support for internal branches of the tree was evaluated by bootstrapping with 10,000 iterations. Tree visualization was conducted using iTOL (version 6.9, 57).

### Phylogenetic analysis with other closely related ST25

Given the close genetic relationships among the study strains, only the first isolated one, F14-11, was used for the selection of closely related strains from other studies. The topgenome (-t) feature of WhatsGNU was used to identify the ten closest strains from a database comprising 4,325 genomes of *A. baumannii* available on GenBank. The database was constructed by downloading all publicly accessible genomes of *A. baumannii* from NCBI in February 2023 (Genome database, search criteria: assembly level = {Chromosome, Complete, Contig}, exclude partial, exclude anomalous).

To expand the diversity of the strains set, the top genomes of strains from the top 10 closest strains of F14-11 as well as four additional ST25 strains (OIFC143, CI79, Naval18 and 107m) were also included. To prevent erroneous conclusions regarding geographic proximity, only one strain per study was included. Incomplete sequenced genomes were excluded. All the genomes were mapped to the F14-11 chromosome set as a reference using the Gubbins script generate_ska_alignment.py, which creates an alignment using SKA2 (version 0.3.7, 58). The resulting alignment measured 4,061,760 bp with 1,511 variable positions. The output was then analyzed by the Gubbins algorithm relying on RAxML using the GTRGAMMA model (version 8.2.12, 58) for constructing the phylogeny in each iteration to generate an unbiased phylogeny. The reproducibility of the node positions in the tree topology was assessed using bootstrap analysis with 10^4^ replicates. No root was assigned because strains known to be outgroups were genetically too distant. Including such distant outgroups would result in insufficient sequence alignment and therefore insufficient information for accurate phylogenetic analysis. Finally, the tree was visualized, incorporating host and isolation location data, using the R package ggtree (version 3.12.0, 58).

### Variant detection analysis

Primary Illumina sequence reads (fastq) of monophyletic isolates were aligned to the earliest (F14-11), using the tool snippy (Galaxy version 4.6.0, 59). The primary Illumina reads of the F14-11 strain were also subjected to the analysis as control and returned no SNPs. Deletions, insertions and rearrangements were identified using the software package MAUVE (version 2.4.0, 60). The list of the identified genetic variations is provided in **TABLE S3**.

### Statistical analysis

All statistical analyses were performed using RStudio 4.1.2 (www.rstudio.com). The medians of the biofilm formation levels of the different strains were compared using the non-parametric Kruskal-Wallis test, followed by a Bonferroni-Holm correction for pairwise comparisons with strain F14-11. The potential correlation between chromosome size and the number of months elapsed between isolation dates was evaluated using Spearman’s correlation test. For months with multiple strains isolated, the mean chromosome size of those strains was used.

### Antibiotic susceptibilities testing

Susceptibility of the strains to a panel of antibiotics was evaluated by disc diffusion on Mueller-Hinton agar (Biorad, France) following the CA-SFM 2013 recommendations (https://resapath.anses.fr/resapath_uploadfiles/files/Documents/2013_CASFM.pdf).

Inhibition values were interpreted according to CA-SFM 2013 breakpoints for all antibiotics but aztreonam, for which *Pseudomonas* spp. breakpoints are given. The strain *Pseudomonas aeruginosa* CIP 7110 was used as control for all the antibiotic susceptibility testing. MIC experiments were also conducted to evaluate susceptibility to colistin and piperacillin according to EUCAST guidelines (37).

### Biofilm formation assay

Biofilm-forming capacities of the isolates were evaluated using the crystal violet staining method, with slight modifications to the protocol previously described (38). Briefly, exponential phase cells were inoculated into 96-well polystyrene plates in 200µL of BM2 minimal medium (39) supplemented with 10mM potassium glutamate (BM2G) at starting OD_600_ of 0.1 and incubated at 37°C in the absence of light for 48 hours, without shaking. The medium was replaced with fresh BM2G medium after 24 hours of incubation. After 48 hours, the cells in suspension were removed and the adherent cells were fixed with 3.7% formaldehyde for 10 minutes. The wells were then washed twice with distilled water and 200µL of 0.1% crystal violet was added to each well, incubated for 10 minutes, and washed three times with distilled water. To solubilize the crystal violet, 200μL of 95% ethanol was added and the absorbance was measured at 590nm with Infinite M200 PRO (TECAN) and Magellan 7.1 SP1 (TECAN).

## Results

### A ST25 monophyletic infection related to human and animal cases

A retrospective study was performed on *A. baumannii* isolates cultured from a single animal-patient (cat) over a 5-year period (2014 to 2019). This patient-animal developed a urinary *A. baumannii* infection following the placement of a urinary invasive medical device. The early isolate was identified as multidrug resistant, including to human-restricted antibiotic carbapenems, but not to colistin. However, two colistin treatments failed to clear the infection. Given the limited antibiotic arsenal authorized in veterinary medicine, and the absence of invalidating clinical signs, no further antibiotic treatment was attempted. During medical follow-ups, eight strains (referred as longitudinal strains) were sequentially isolated and stored by the hospital laboratory from November 2014 to February 2019 (**FIG S1 & TABLE S1**). In addition, nine strains (referred to as synchronic strains) were subsequently isolated from a single complete urine sample collected on October 2019.

Whole-genome sequencing and hybrid assembly of the seventeen study strains revealed clonally related strains belonging to sequence type (ST) 25 clone, ST associated with UTI in human and animals (6, 20). To identify the closest genomes to F14-11, the earliest isolate, we used the Similar Genome Finder utility of the WhatsGNU tool to query an *A. baumannii* genome database retrieved from the NCBI (5234 genomes, downloaded on February, 17^th^ 2023, 23). Most strains were from human origin as publicly available *A. baumannii* genomes are mostly related to human medicine. The phylogenetic tree constructed from twenty-three ST25 strains reveals an apparent correlation between isolation location and phylogenetic proximity for Eurasian and American distributions. However, both North American strains were potentially military imports from overseas explaining their clustering with an Iraqi strain (41) (**FIG 1**). This analysis also indicated a clustering of clones independently of their host implying zoonotic transfer and highlighting the relevance of investigating animal infections for human health (42). The closest human strain, 16A1524, was indeed isolated in France in 2016. Interestingly, the closest strain, 40288, was also isolated in France from a dog UTI in 2015 (30). F14-11 and 40288 exhibit only 31 SNPs across their chromosomes and 1 SNP between their ∼145Kb-length plasmids p40288 and pF14-11 (**TABLE S2**). The pF14-11 backbone was also detected in 22 of closest genomes (coverage from 65% to 100%, **TABLE S2**). Despite its reduced size, pF14-11 shares 100% identity with larger plasmids pVB82 (215Kb) and pHWBA8 (195Kb) carried by close eponymous strains (**FIG S2** and **TABLE S2**). This size reduction may explain the inability of pF14-11 to undergo conjugation (**FIG S2**).

**FIG 1.**
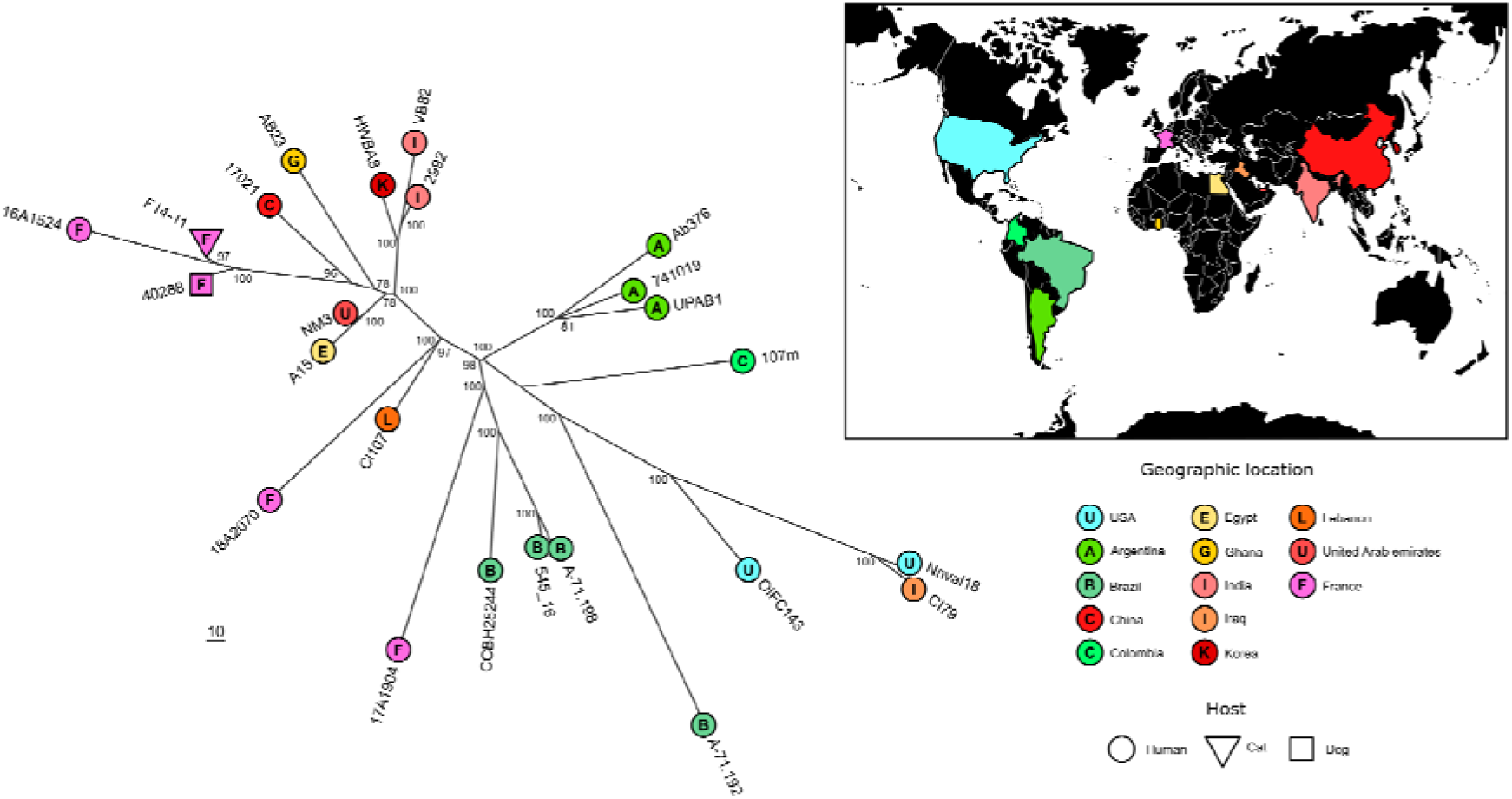
Phylogenetic proximity of F14-11 to other strains isolated from human and animal infections worldwide. (A) Unrooted Maximum Likelihood phylogeny constructed using Gubbins based on RAxML. Tips are colored by geographic origin, labeled with the country’s initial, and shaped by host type. Bootstrap values are shown at nodes. Scale indicates branch length in base substitutions. (B) World map with countries highlighted according to the specified color scheme.

### Genome reduction and insertion sequences dynamics

To further investigate within-host strain diversity, we performed whole-genome comparison of the sixteen study strains to the early one F14-11, set as a reference (**FIG S1** & **TABLE S1**). This analysis revealed a low number of SNPs scattered throughout the genome in comparison to F14-11, with a mean of 31 SNPs per genome ranging from 3 SNPs for the closest genomes to 78 SNPs for the most distant one (**TABLE S1**).

Phylogenetic analysis suggests an absence of correlation between isolation date and phylogenetic relationships (**FIG 2**). Notably, longitudinal isolates F18-02 and F19-02 form an independent clade, referred as clade I, but sharing a hypothetical common ancestor with all other strains. In contrast, late synchronic strains isolated in 2019 cluster with strains isolated in 2015 implying the coexistence of different lineages during the infection. A correlation is observed between the chromosome size of strains and the time elapsed (R²=0.86, P=6.7×10^-3^, Spearman test) indicating a reduction in size during infection (**TABLE S1**). Genome reduction operates on MGE such as prophages. Indeed, most strains show independent excisions of one to three of the four prophages identified in strain F14-11. Close strains F18-02 and F19-02 lost one and two prophages respectively (**FIG 2**, large deletions highlighted in green and **FIG S3A**, prophage regions 2 and 4). The endogenous plasmid underwent further size reduction with four of the synchronic isolates lack a 12-18 kb plasmid region corresponding to the BREX phage resistance system while three have lost the plasmid entirely (**FIG 2 & S3B and C**) (43).

**FIG 2.**
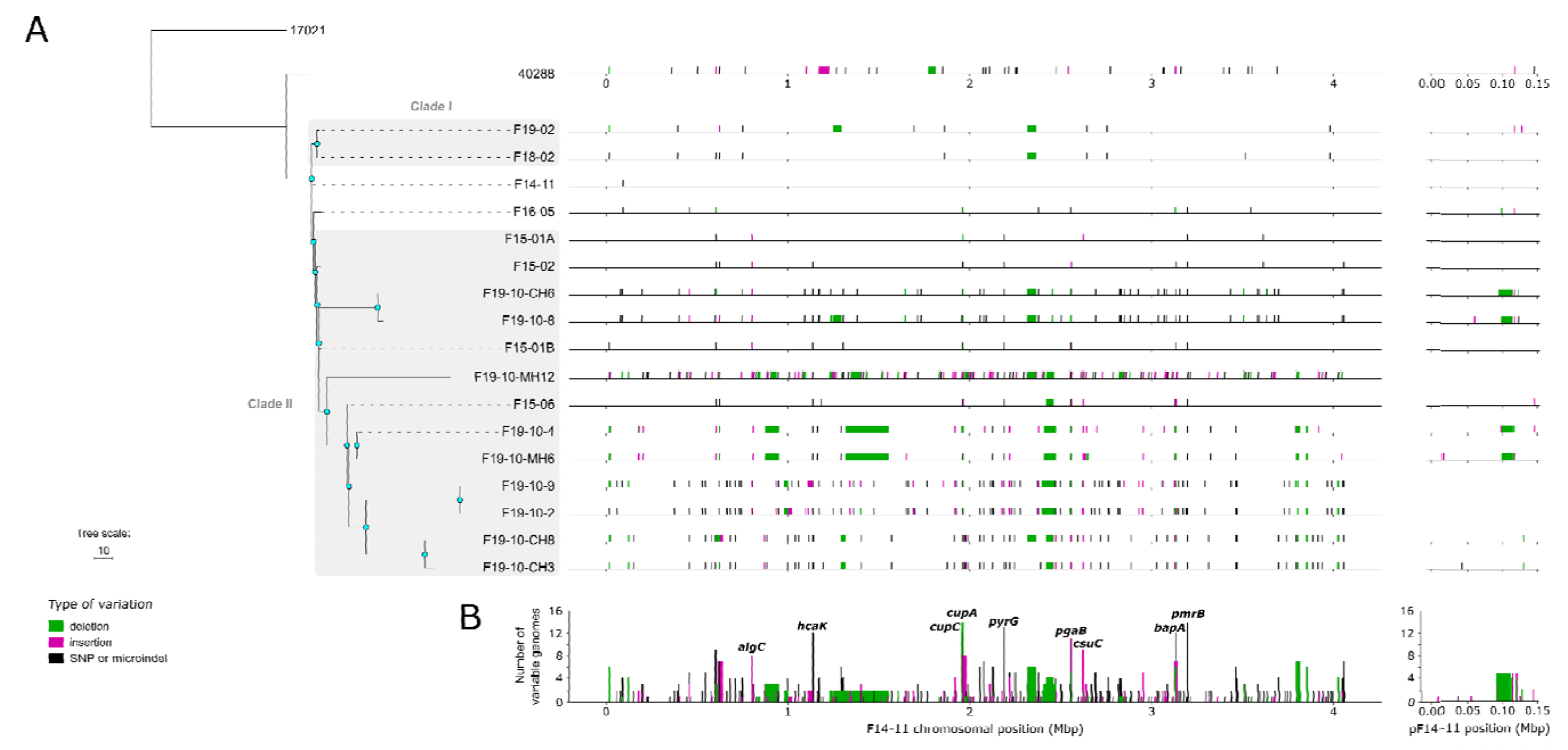
Phylogenic tree of the study strains and 40288 associated with the distribution of the genetic variations. (A) Left: Maximum Likelihood ylogeny constructed with Gubbins based on RAxML. The tree was rooted with strain 17021. Scale indicates branch length in base substitutions. e nodes indicate bootstrap values >80. Right: Representation of SNPs and microindels in black, insertions in pink and deletions in green along the nomes in comparison to F14-11 (reference genome). (B) Cumulative distribution of the genetic variations along the F14-11 genome with *y*-axis icating the number of study strains harboring the genetic variation. The genes most prone to variation are shown.

Insertion sequences (IS) recombination led to chromosome deletions. The most striking evidence of this phenomenon is the deletion of a 36.6kb region, fixed in a subclade of eight strains within clade II, flanked by to identical IS*Aba34* in F14-11 (**FIG 2** and **FIG S3**). In addition, in comparison to F14-11 chromosome, we observed multiple new chromosomal insertions by an IS originated from the plasmid (**FIG S4**). Indeed, IS*17*, an IS*5* family insertion sequence, was absent from the chromosome of the earliest strain, F14-11, but carried by its plasmid pF14-11. While a single instance of IS*17* was found in the chromosomes of strains isolated in 2015, it significantly multiplied in synchronic strains, climaxing with 75 occurrences in strain F19-10-MH12. Interestingly, IS*17* is also present on p40288 of related 40288 strain although retrained to this replicon.

### Recurrent variation of adhesion and biofilm genes

Over 82% of SNPs and microindels of the strains studied are in coding sequences, with 80% of non-synonymous SNPs. Several of these genes are involved in adhesion and biofilm formation – genes that are also affected by insertion and deletion events (**FIG 2**; *cup* (44), *csu* (45), *pga* (46), *bapA* (47), *algC* (48) genes). All clade II strains exhibit at least one mutation in one of these genes, while clade I strains retain genetic sequences identical to those of F14-11. Strikingly, genes involved in biogenesis of archaic and classical types of chaperone-usher pili involved in adhesion step of biofilm formation (*csu* and *cup* operon respectively) were subjected to IS insertion, deletion and frameshift. Specifically, the *csu* operon is affected by insertions by distinct IS in strains F15-01A (IS*17*) and F15-06 (IS*Aba33*), this latter insertion being conserved in most of synchronic isolates (7 out of 9). Genes of the classical pili Abp1 (formerly Cup), *cupA* and *cupC*, were also affected by large deletions and indel leading to frameshift in most strains of the clade II. Additionally, genes involved in biofilm formation such as *algC* and *pgaABCD* but also *bapA* gene were also disrupted by IS insertion or subjected to deletion events (**FIG 2**).

Hence, during the infection, may co-exist strains with varying adhesion and biofilm abilities. To bring experimental evidence to this hypothesis, biofilm-forming capacities of the study strains were measured and compared with those of the environmental DSM30011 strain described as strong biofilm-producer (**FIG 3**) (39). As observed for human clinical strains, the animal clinical strain F14-11 forms less biofilm than DSM30011 (3-fold; p-value: 0.00143). However, F14-11 carries a plasmid closely related to plasmid pAB5 that downregulates biofilm *pga* and adhesion *cup* genes in strain UPAB1 (20, 21). We therefore tested the hypothesis that pF14-11 may restraint biofilm production. Yet, pF14-11 curing did not increase adhesion, biofilm nor PNAG production (**FIG S5**). We also tested if the residual biofilm capacity of F14-11 could involve type IV pili or *prp* type I pili but fail to identify its adhesion mechanism (**FIG S5**).

**FIG 3.**
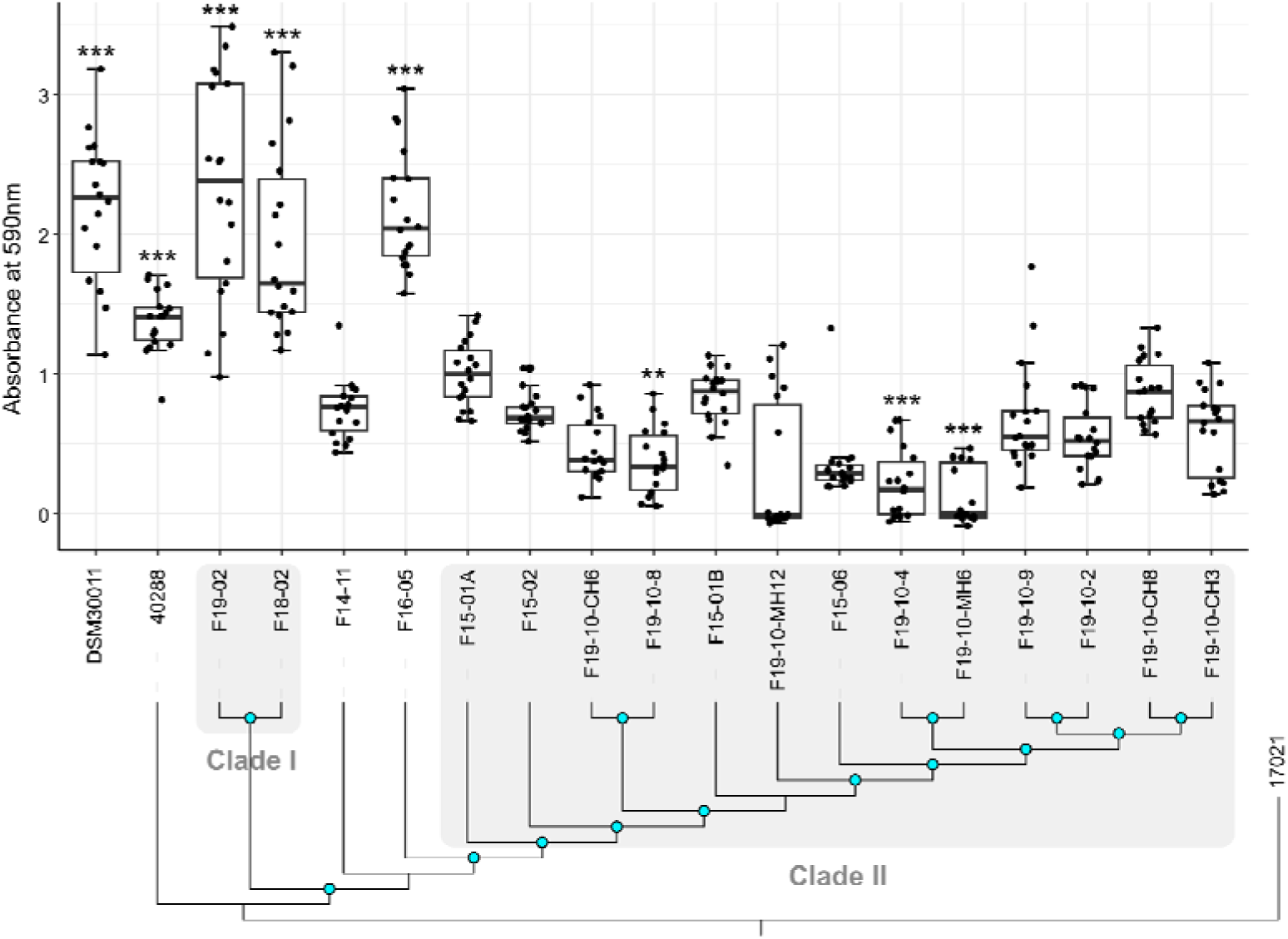
Biofilm formation by the study strains and the dog UTI strain 40288 associated with their phylogenetic relationships. Blue nodes indicate bootstrap values >80. Biofilm formation was assessed by measuring the optical density of crystal violet stained matrix and compared to environmental strain DSM30011. Error bars are generated from 12 replicas from 4 independent experiments. Statistically significant differences with F14-11 (Kruskal-Wallis rank-sum tests with Bonferroni-Holm correction) are highlighted with asterisks: (*) if the p-value < 0.05, (* *) if the p-value ≤ 0.01, and (* * *) if the p-value ≤ 0.001.

Surprisingly, F16-05 isolate is a strong biofilm producer despite a frameshift in the *cupC* gene and a non-synonymous SNP in *pgaB*. However, its adhesion capacity is indeed greatly affected (**FIG S5**). Moreover, in spite of the genome proximity between F14-11, F18-02 and F19-02 these latter two strains produce biofilm amount comparable to DSM30011 (p-value=1) (**FIG 3**). However, none of the SNPs, mainly found in intergenic region, nor the phage deletions of these strains are common between F16-05 and the late isolates and could account for this phenotype. One hypothesis would be that biofilm production is repressed in F14-11 strain by a chromosomally encoded factor. In contrast, strains from the clade II exhibit comparable or reduced biofilm production in comparison to F14-11. However, considering the numerous adverse variations in their adhesion and biofilm genes, a more drastic reduction in biofilm formation was expected. In addition, F14-11, despite identical biofilm and adhesion genes (*cup*, *csu*, *pga*, *algC* genes), produces less biofilm than dog UTI strain 40288 (p-value: 1.3e-05). These results indicate that the F14-11 strain isolated at an early stage of infection exhibits low biofilm production, possibly due to repression of biofilm genes by a chromosomally encoded factor. These genes were subjected to recurrent mutation in subsequently isolated strains.

### Limited conserved metabolism modification

Bacterial metabolism versatility is a crucial factor for successful colonization of a new environment. Similarly to humans, cat’s urine composition is highly variable and complex with thousands of unique compounds (49, 50). The variations occur both among different individuals and within the same individual over time, under the influence of environmental factors, particularly the type of diet (49). Interestingly, genetic variations in study strains mainly occur in metabolic genes (**FIG S8**).

However, only two metabolic entities appear recurrently affected in the isolated strains: the *pyrG* gene and the *hca* operon. Most strains (13/17) have a substitution in *pyrG* gene encoding the putative PyrG enzyme catalyzing amination of UTP to CTP thereby regulating intracellular CTP levels (**FIG 2**) (51). The non-synonymous substitution (H193R) being in the vicinity of the conserved putative UTP binding site we hypothesized that it increases the enzyme affinity for its substrate. We also observed repeated gene variation (for 14 strains out of 17) for aminobenzoate utilization (*hca* operon, **FIG 2**). Strains presented either a non-synonymous substitution (G269V) in the putative membrane transporter HcaK, or a stop codon mutation in the gene encoding its putative repressor HcaR, indicating a possible increase in aminobenzoate utilization. This family of compounds can be found in human and animal food, sometime used as food preservative, excreted in urine (52). Metabolic capacities may also be affected by deletion events. For instance, the 36.6kb deletion beforementioned, encompasses 27 genes including 19 metabolism genes whereas only 40.8% of F14-11 genes are of this category (1533/3758). This conserved large deletion suggests a dispensability of associated metabolic functions for growth in bladder.

We further explored the metabolism capacities of study strains using Biolog metabolic phenotypic microarray (**FIG 4**). Overall strains demonstrated a poor ability to grow in carbohydrates but were able to utilize most amino acids as sole nutrient source. This shared metabolic profile is advantageous in the context of urinary infection as urine usually lacks carbohydrates whereas amino acids and small peptides are abundant (49, 50). The metabolic profiles of the different strains were mostly similar, with the exception of F19-10-MH12 strain whose metabolic capacities may have been profoundly affected by numerous IS*17* insertions (**FIG 4 & S4**). For the set of nutrient sources tested, metabolic capacities were inconsistently altered and could be explained by allelic variation and transposition. For instance, a P51H substitution likely abrogated GABA permease function resulting in a growth defect in GABA in strains F19-10-8 and F19-10-CH6. IS*Aba53* insertion upstream an amino acid transporter and downstream an arginine *N*-succinyltransferase may account for slower growth on L-arginine, with ∼2-fold lower area under the curve over 48 hours (AUC_48h_) of strains F19-10-2 and F19-10-9 compared to F14-11. More surprisingly, seven strains exhibit a more efficient growth with D-malic acid with up to ∼5-fold higher AUC_48h_ compared to F14-11 (F19-10-2, F19-10-9, F19-10-CH3, F19-10-CH6 and F19-10-CH8). This gain of function for utilization of an artificial amino-acid may reflect an increase efficiency of enzymes from the tricarboxylic acid (TCA) cycle and the requirement of this metabolism pathway for growth in the urinary tract as observed for Uropathogenic *E. coli* (53). An alternative explanation would be an increased metabolization of D-malic acid that is a food preservative authorized in animal feed (54). This unexpected gain in function, may therefore reflect adaptation to the host’s urine composition.

**FIG 4.**
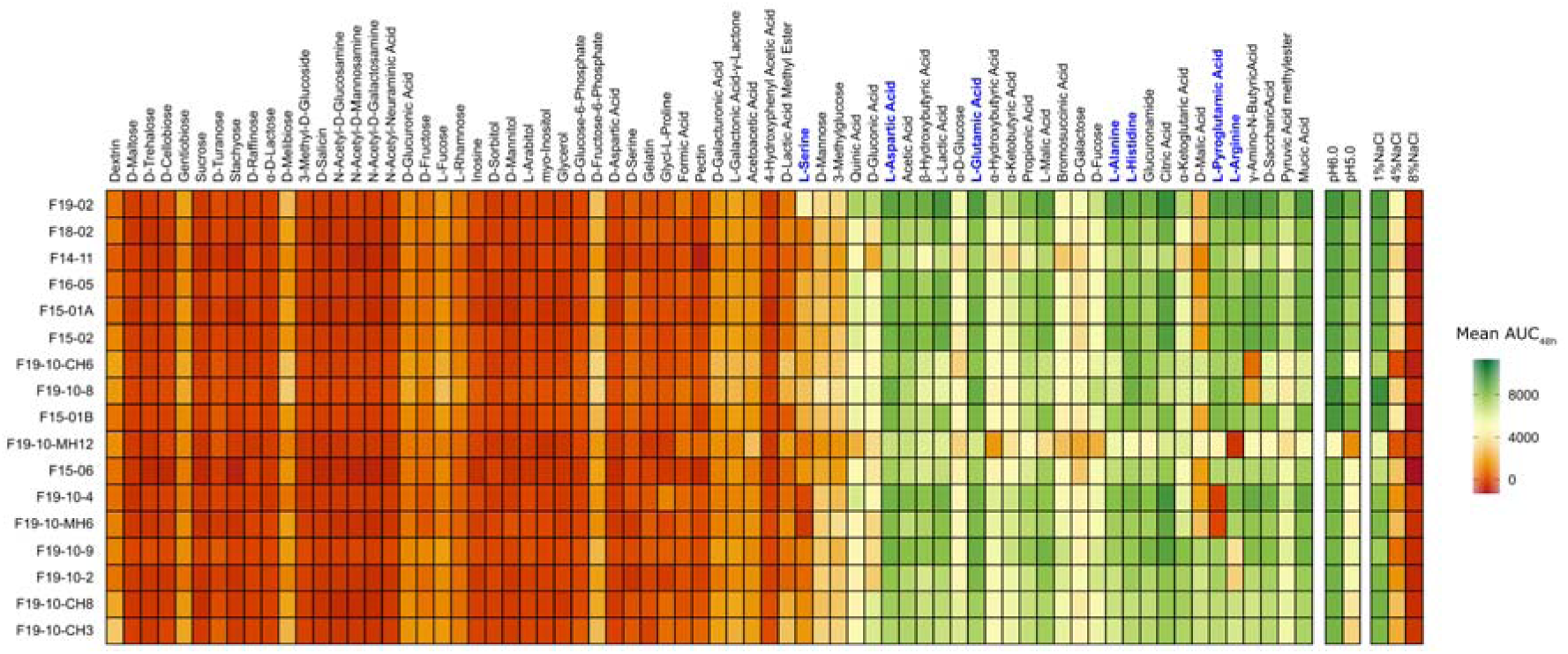
Heatmap of AUC_48h_ for growth either in 70 different nutrient sources, or two pH and three salinity conditions. Amino acids are highlighted in bold blue. The AUC_48h_ parameter was extracted from kinetic data over 48 h in Biolog Gen III MicroPlate. The highest and lowest AUC_48h_ are indicated by green and red colors, respectively. The x-axis lists the tested compounds and conditions, while the y-axis lists the study strains, which are displayed in the order corresponding to the phylogenetic tree presented in Figure 2A. Mean data of two independent experiments are shown.

### Overall resistance loss contrast with colistin resistance fixation

The first isolate F14-11 is resistant to penicillins, cephalosporines (but ceftazidime), fluoroquinolones, aminoglycosides (but amikacin) and carbapenems. Following colistin treatments, colistin resistance developed in 2015 isolates (**FIG 5**). Colistin resistance was mainly maintained after cessation of treatment with the exception of two sensitive isolates (F18-02 and F19-02). Colistin resistant isolates share a PmrB(P170T) substitution. We ensured that PmrB(P170T) transfer to F14-11 resulted in colistin resistance (MIC>16µg/mL). Contrarily to PmrB(P170L) associated to a growth defect in laboratory define medium (55), F14-11 derivatives PmrB(P170T) and parental F14-11 have comparable growth rate (**FIG S6**). Isolation of colistin-sensitive strains with wild type *pmrB* sequence suggests coexistence of both resistant and sensitive strains (**FIG 5** ; F18-02 and F19-02).

**FIG 5.**
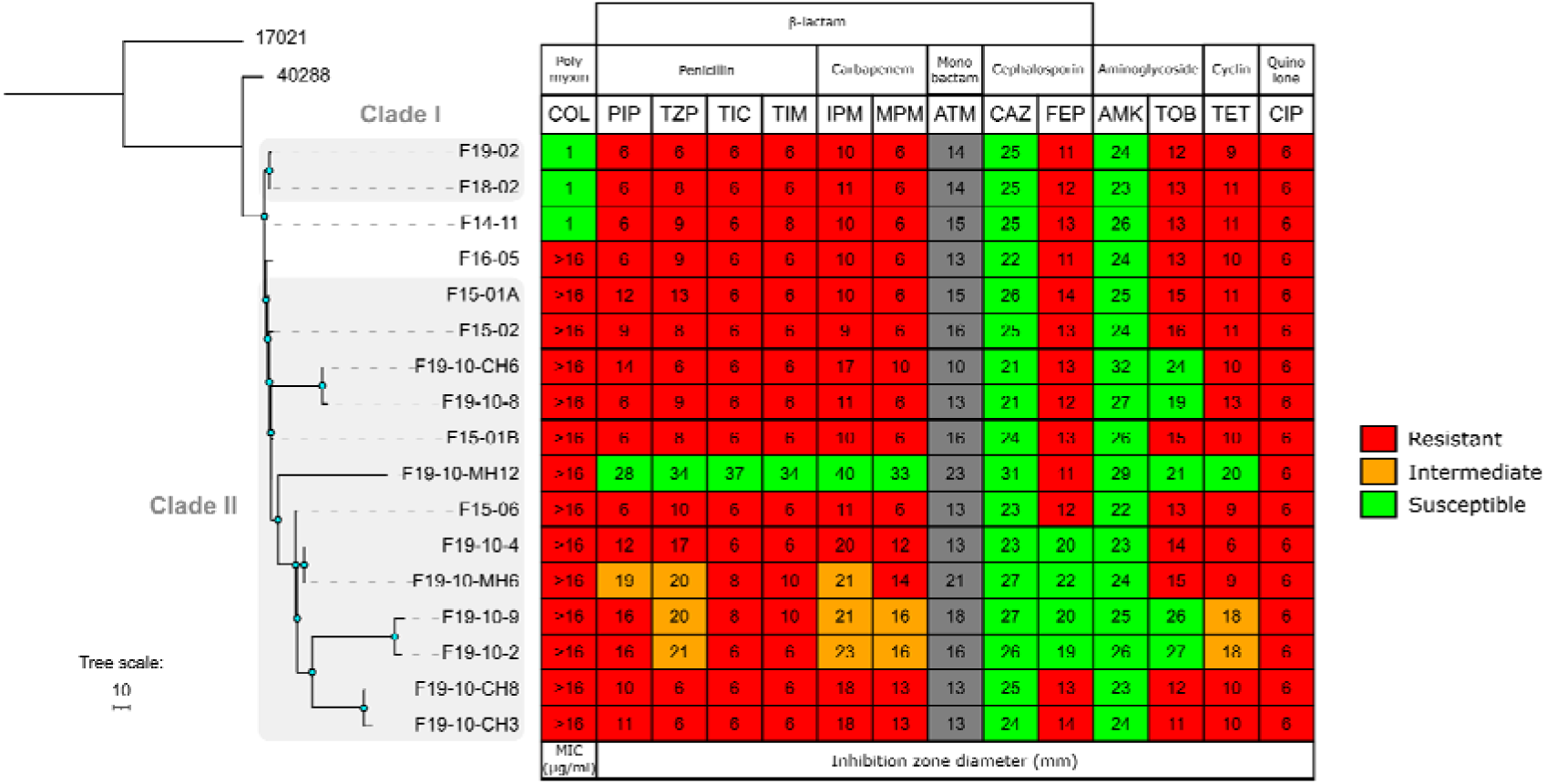
Heatmap of antibiotic susceptibility of the study strains associated with their phylogenetic relationships. Blue nodes indicate bootstrap values >80.. Abbreviations. AMK: Amikacin, GEN: Gentamicin, TOB: Tobramycin, CIP: Ciprofloxacin, PIP: Piperacillin, TZP: Piperacillin + tazobactam, TIC: Ticarcillin, TIM: TIC + clavulanic acid, CAZ: Ceftazidime, FEP: Cefepime, IMP: Imipenem, MEM: Meropenem, ATM: Aztreonam, TET: Tetracycline, COL: colistin. Antibiotic susceptibilities were either measured using minimal inhibition concentration (MIC) or by the disk diffusion methods. The antibiotics tested are listed on the x-axis and the study strains on the y-axis.

In contrast to colistin resistance, several antibiotic resistances were lost in the clade II in comparison to F14-11. Most susceptibilities emerged due to plasmid loss. Early isolate plasmid pF14-11 harbors genes conferring resistance to tetracycline (*tetA(B*)), aminoglycosides (*strA/strB, aac*(*3*)*-IIa, aac(6’)-Ian*) and sulfonamides (*sul2*) frequently associated with this family of plasmid (42). Plasmid loss expectedly results in susceptibility to tetracycline and tobramycin as observed for strains F19-10-MH12, F19-10-2, and F19-10-9 (**FIG 5**). However, plasmid loss alone may not explain recovered susceptibility to antibiotics as two tobramycin-sensitive strains still carry a plasmid (**FIG 2 & 5**; F19-10-CH6 and F19-10-8). Their intact resistance gene sequences led us to hypothesize that plasmid-specific deletion events encompassing a putative DNA primase encoding gene may lower plasmid copy number and maybe decrease associated resistance to tobramycin.

Chromosomal resistance carried by mobile genetic elements were also subjected to curing. F14-11 strain carries two copies of Tn*2006* carrying *bla_OXA-23_* gene accounting for its carbapenem resistance. Closely related strain 40288 also bears two Tn*2006* copies located either in an AbaR4 resistance island inserted in the *comM* gene or interrupting the *vgrG* gene (30) . Compared to 40288, F14-11 present a recombination event between Tn*2006* copies leading to a split AbaR4. Further deletions of one or two Tn*2006* in several strains may explain increased susceptibility to carbapenems (**FIG 5** ; respectively in F19-10-4, F19-10-CH3, F19-10-CH8 F19-10-MH6 and F19-10-MH12). Additionally, resistance decreases without antibiotic gene loss may be related to compromised growth in laboratory medium associated with large chromosomal deletion events (**FIG 5 & S7**; F19-10-2, F19-10-9 and F19-10-MH6). Taken together, these results indicate an overall decline in antibiotic resistance mediated by curing of mobile genetic elements. They also suggest coexistence of strains with varying resistance levels within a population hence illustrating within host heteroresistance in the absence of antibiotic pressure.

## Discussion

High versality of certain clones of *A. baumannii* may explain their ability to colonize various host and dominate the global epidemiology of this pathogen. In this study, we examined within-host genomic evolution during prolonged urinary infection of a ST25 strain in an animal host. The genetic proximity between this animal strain and human strains strongly favors the hypothesis of a zoonotic transmission, with a potential transmission from a human to animal hosts (a cat for F14-11 strain and a dog for 40288 strain). A relatedness between human and animal isolates of the ST25 lineage was also suggested by a recent study (42). Remarkably, most animal ST25 strains analyzed in this latter study were isolated from urinary infection in greater extent than for human strains suggesting a host-specific tropism. Following transmission, strains may further evolve by mutations, insertions and deletions presumably mediated by MGE-driven intragenomic rearrangements, particularly through IS dynamics known to promote evolution *in vitro* (56, 57). In our study, the single plasmid copy of IS17 was subsequently detected in the chromosomes of strains collected during infection and extensively propagated in the genomes of late isolates, leading to the pseudogenization of accessory genes. Additionally, recombination of IS*Aba34* led to the loss of a large chromosomal fragment encompassing genes mainly involved in metabolism. While IS element expansion and subsequent recombination events are traditionally associated with the restricted lifestyle of endosymbionts and their dependence on the host, genome reduction have recently been linked with pathogenicity (58). These results bring additional evidence of IS dynamics shaping bacterial evolution to adapt to a new environment, as recently described for the plasmid-mediated IS*1* transposition in clinical Enterobacteriacea (59).

Genetic variations are mainly observed in genes associated with biofilm formation and metabolism. Strikingly, we found that two of three type I *pili* structures encoded by these ST25 isolates were profoundly affected by mutations (*csu* and *cup* genes). Interestingly, several studies described recurrent downregulation or mutation affecting of the *csu* operon following infection by human clinical isolates (15, 55, 60). Besides type I *pili*, other biofilm formation genes present genetic variations such as *algC* and *pga* operon. The *cup* and *pga* genes were also described as downregulated by large conjugative plasmid of a urinary strain involving plasmid-encoded transcriptional regulators that directly or indirectly repressed their expression (20). For the present study urinary strain, curing the related large plasmid did not modify adhesion pattern suggesting that a strong repression of these genes may also involve chromosomal determinants. Repression of biofilm and adhesion gene expression indicates a dispensability for infection and led to their evolution by mutation. This hypothesis is strongly sustained by the recurrent and varying types of gene inactivation observed in those loci. Adhesive structures such as *pili* are highly immunogenic and are even exploited for vaccine development (61). Loss of these appendages would favor immune escape.

Furthermore, our data suggest a coexistence of strong and poor biofilm-producers rather than a homogenous population. This coexistence was also described in other urinary pathogen during a long-term human urinary infection with *Klebsiella pneumoniae* (62). These observations may indicate heterogeneous occupation of the urinary tract with spatially distinct niches. Alternatively, strong and poor biofilm-producers could collaborate, with sufficient extracellular matrix to allow poor biofilm-producer strains to embed themselves in the biofilm. The metabolic genes have undergone the strongest evolution within the host during this long-term infection. This trend is also observed in longitudinal within-host studies conducted on other bacterial species indicating common survival strategies (63–66). However, strains display an unexpected stable metabolism profile. This observation may reflect our technical limitation as only 70 nutrient sources were tested. Nevertheless, the reduced genomes may be sufficient to efficiently grow on nutrients available in their host environment, such as amino acids, and even to adapt to this specific medium with as near neutral pH, low sodium and chloride, and even presence of diet-associated compound (D-malic acid metabolism). This latter observation brings new perspectives to the management of patients suffering from *A. baumannii* infection, and the potential role of host diet in patient care. Our study also present divergences with previous studies on human infections. Indeed, we observed only few mutations in translational machinery genes or in genes related to replication, recombination, and repair (16, 60). However, these evolutions may have been shaped by alternative antibiotic treatments performed on human patients. Moreover, contrarily to previous report, colistin resistance was maintained in the study strains. Snitkin *et al.* isolated colistin-sensitive strains from human patients after removal of this antibiotic (60). However, this study used a single-isolate approach randomly isolated that may not reflect the overall population. Alternatively, the PmrB(P170T) described in the present study may represent a low-cost mutation or even be beneficial in the context of urine infection. Indeed, recent findings on PmrAB indicate that this two-component system responds to low pH (5.8) and metal ions (Fe^2+^, Zn^2+^ and Al^3+^) (67). Therefore, the PmrB(P170T) mutation could have been maintained to mitigate a urine component toxicity.

Our results also unveiled a hidden phenotypic and genetic diversity within a clinical sample. Rather than to restrict conclusion to a single isolate with defined characteristics, the advent of new sequencing technology such as single-cell sequencing should enable a comprehensive understanding of pathogen genomic dynamics during infection (68).

## Acknowledgements

We thank the HORIGENE group members for useful comments throughout the study. We are grateful to Michel Pépin and Brigitte Chataigner from Laboratoire Vétérinaire Départemental (VetAgro Sup) for allowing access to *A. baumannii* animal isolates. We thank Agnese Lupo and Marisa Haenni (Anses Lyon, Unité Antibiorésistance et Virulence Bactériennes, France) for performing antimicrobial susceptibility tests and Maxime Bruto (Anses Lyon, UMR Mycoplasmes, France), Lucie Etienne and Carine Rey (CIRI, France) for fruitful discussions and comments on phylogenetic analysis.

