## Supplemental methods and figures for "Phenotypic and genetic heterogeneity of *Acinetobacter baumannii* in the course of a chronic infection"

### Supplementary materials

 Authors: Léa Bednarczuk^a,b^, Alexandre Chassard^a#^, Julie Plantade^a#^, Xavier Charpentier^a^ and Maria-Halima Laaberki^a,b*^.

a. CIRI, Centre International de Recherche en Infectiologie, Inserm, U1111, Université Claude Bernard Lyon 1, CNRS, UMR5308, École Normale Supérieure de Lyon, Univ Lyon, 69007, Lyon, France

b. Université de Lyon, VetAgro Sup, 69280 Marcy l'Etoile, France.

\# equivalent contributions

**Supplementary methods**

**Supplementary figures**

**Supplementary methods**

**Conjugation assays**

Rifampicin-resistant derivatives *A. nosocomialis* M2 and *A. baumannii* strains AB5075 and A118 (1) were used as recipient strains for plasmid donor strains (ATCC17978 carrying pAB3 or F14-11 carrying pF14-11). Overnight cultures in liquid LB were grown at 37°C for recipient and donor strains were mixed equally and 50 µl aliquots were spotted on LB agar medium. Each culture was also mixed with the same volume of LB medium for control spots. After 24h incubation at 37°C, bacteria were resuspended in LB medium and plated on selective medium (rifampicin 100 µg/ml in combination with tetracycline 30 µg/ml for conjugation of pF14-11, or with sulfamethoxazole 30 µg/ml and trimethoprim 5 µg/ml for conjugation of pAB3), as well as on LB medium supplemented with rifampicin (100 µg/ml) for numbering of recipient bacteria. Conjugation frequencies were calculated as the ratio of the number of conjugants to the total number of bacteria.

#### Congo red binding quantification

Congo red binding capacities of strains were quantified using the protocol previously described (2). Briefly, two drops of overnight cultures of each strain were incubated on YESCA agar media supplemented with 50 µg/ml Congo red and Brilliant Blue G 1µg/ml at 37°C in the absence of light for 48 hours, without shaking. The cells were then recovered and washed twice with 50 mM potassium phosphate buffer. The cells were resuspended in 1 mL of this buffer and the OD_600_ was adjusted to 1. A 100 µL aliquot of each sample was transferred to a 96-well polystyrene plates, and the fluorescence of Congo red was measured using a microplate reader with an excitation wavelength of 485 nm and an emission wavelength of 612 nm. Buffer was used as the blank.

#### Adhesion assays

The *sf-gfp* gene was integrated into the *attTn7* site of all strains of interest. Exponential phase cells of the resulting fluorescent strains were inoculated into uncoated 24-well Ibidi microscopy plate in 500µl of LB medium at starting OD_600_ of 0.1 and incubated at 37°C in the absence of light for 1 hour, without shaking. After incubation, the cells in suspension were removed and the adherent cells were fixed with Antigenfix for 15 minutes. The wells were then washed twice with distilled water. Four images per wells were collected with a Yokogawa CQ1 using a 40× objective. Fluorescent bacteria on these images were counted using the Yokogawa data analysis software. This procedure was repeated across three independent experiments. The median number of adherent cells per mm² was compared pairwise using the non-parametric Kruskal-Wallis test, followed by Bonferroni-Holm correction for multiple comparisons, conducted in RStudio 4.1.2 ([www.rstudio.com](http://www.rstudio.com))

#### Growth analysis

Strains were streaked on LB-agar plates and incubated overnight at 37°C. A single colony was then inoculated into 2 ml of the tested medium (LB or BM2G) and incubated overnight at 37°C. The optical density at 600 nm (OD600_nm_) of these overnight cultures was measured and adjusted to 0.001 by diluting the cultures in fresh tested medium (LB or BM2G). The diluted cultures were then transferred to a flat-bottom, transparent 96-well microplate for growth curve analysis. Incubation was carried out at 37°C for 18 hours in LB and 25 hours in BM2G in an Infinite M200PRO TECAN plate reader, with agitation (amplitude: 5 mm) for one minute every 10 minutes, followed by OD_600nm_ measurements. This procedure was performed in three independent experiments, with triplicates for each strain in each experiment.

**Supplementary figures**

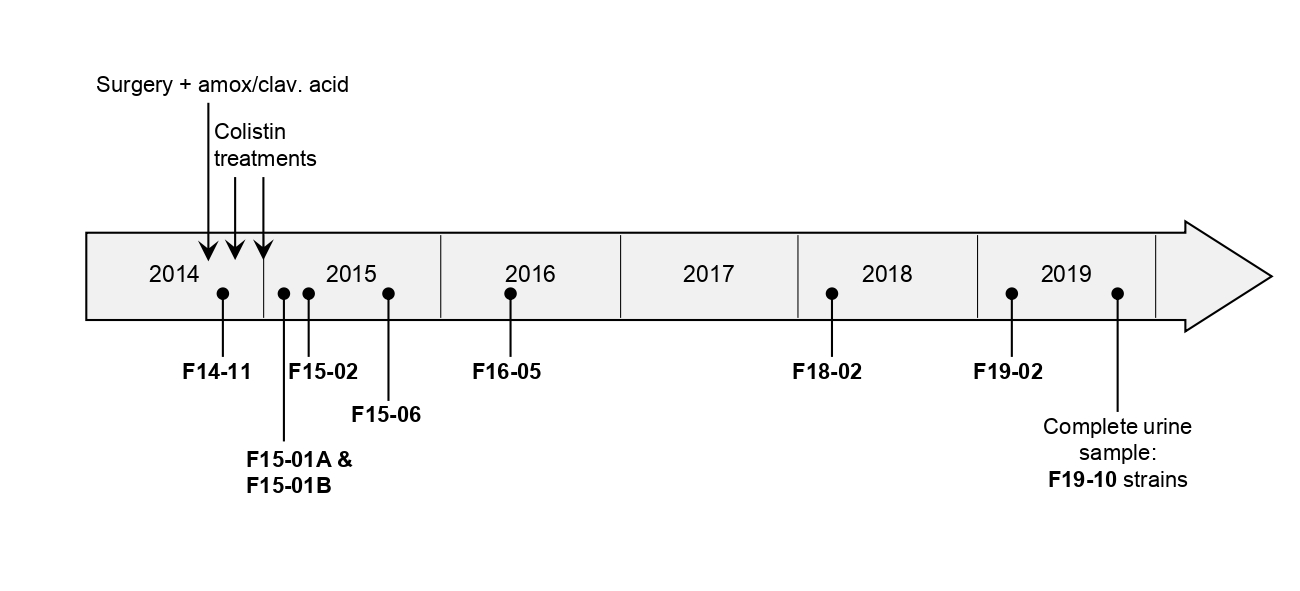

**FIG S1**. Chronological representation of antibiotic treatments and bacterial isolation. Vertical arrows indicate a postoperative antibiotic prophylaxis with amoxicillin-clavulanic acid (amox/clav.) and two colistin treatments. Isolates are named after the year and month of isolation.

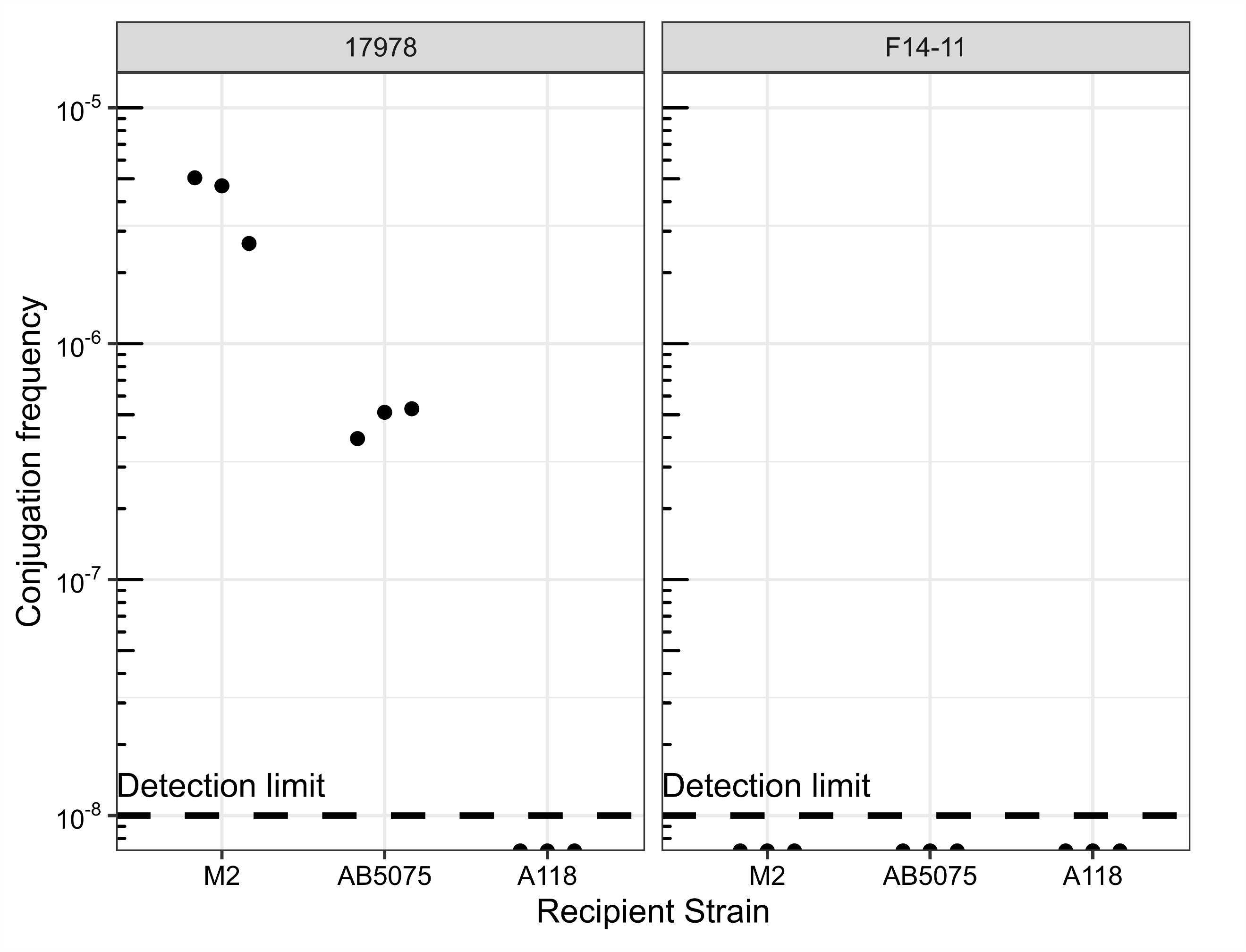

**FIG S2** Conjugation frequencies between (left panel) *A. baumannii* strain 17978, carrying pAB3, or (right panel) strain F14-11, carrying pF14-11, and the recipient strains *Acinetobacter nosocomialis* M2, *A. baumannii* AB5075, or A118. Conjugation of pAB3 was used as control of conjugation conditions.

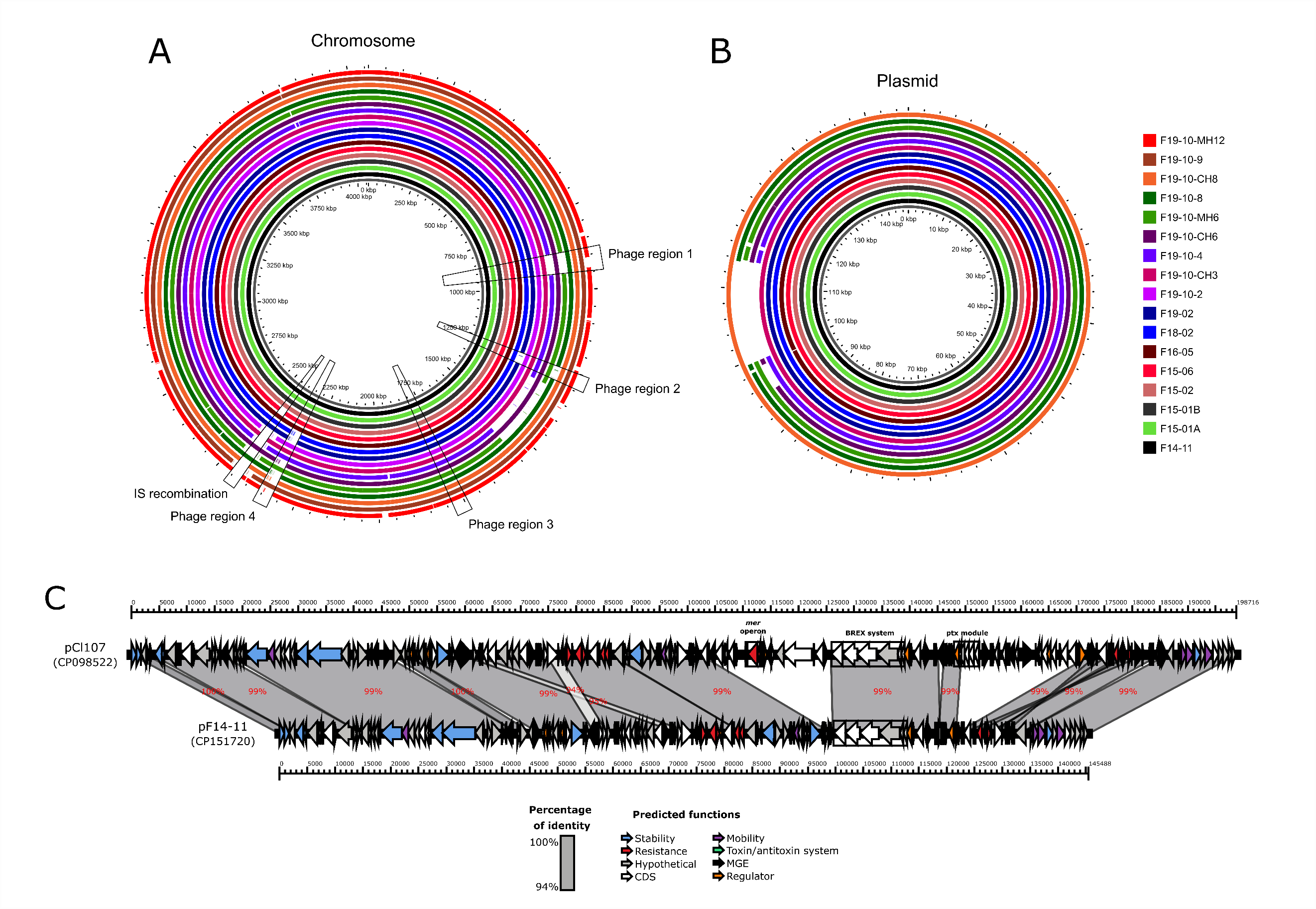

**FIG S3** (A) Chromosome and (B) plasmid maps of the study strains using Gview. (C) Comparison of pF14-11 plasmid with pCl107 from Cl107 strain with Genofig revealing conserved and varying features with this well-described plasmid (3). Schematic representations are drawn to scale. Arrows represent open reading frames and prediction functions relevant to this study are indicated in the legend.

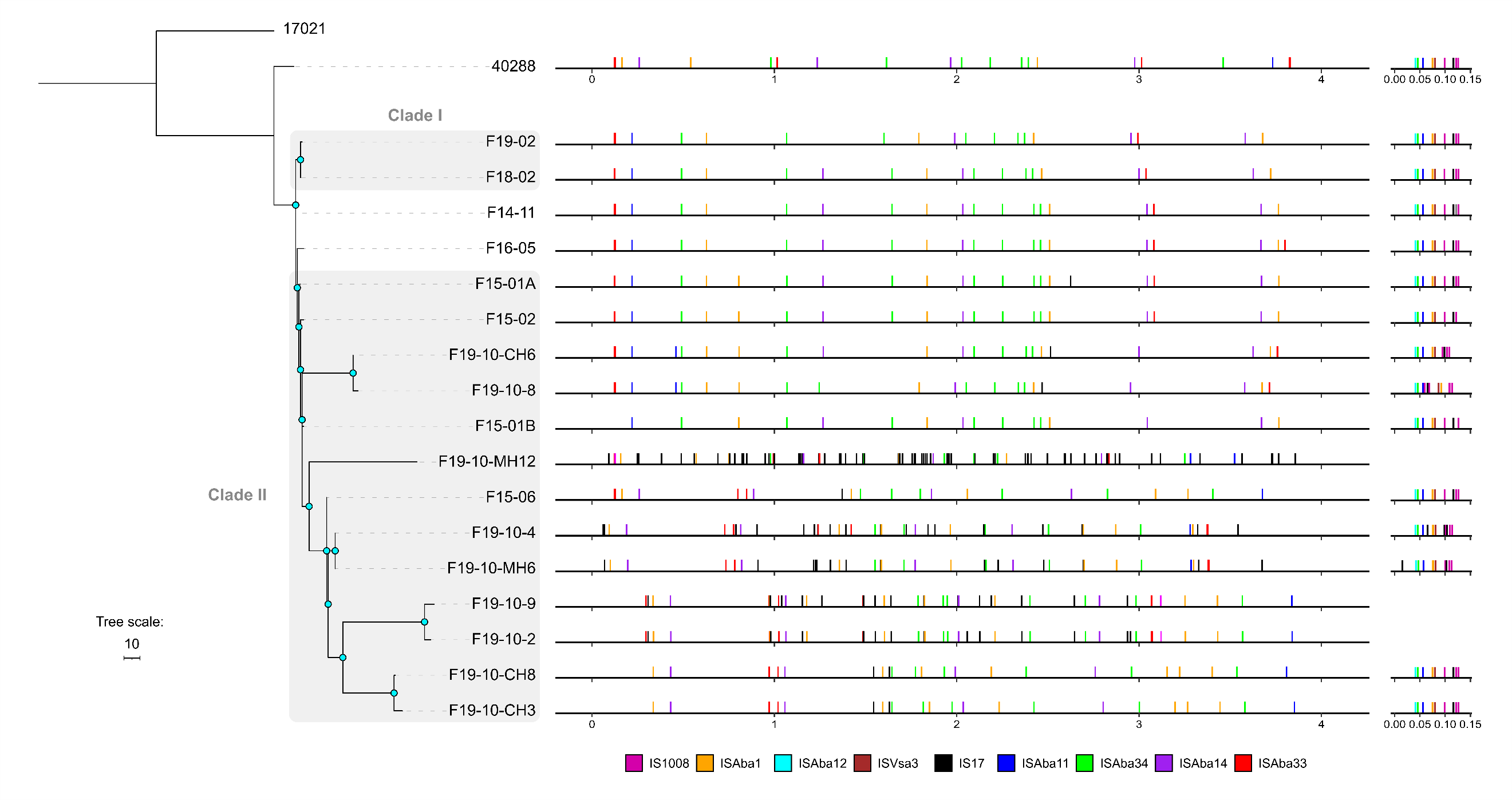

**Figure S4** Distribution of insertion sequences identified with ISfinder (4) in study strains and 40288 strain along their respective genomes associated with the phylogenetic relationships. Blue nodes indicate bootstrap values >80. Scale indicates branch length in base substitutions.

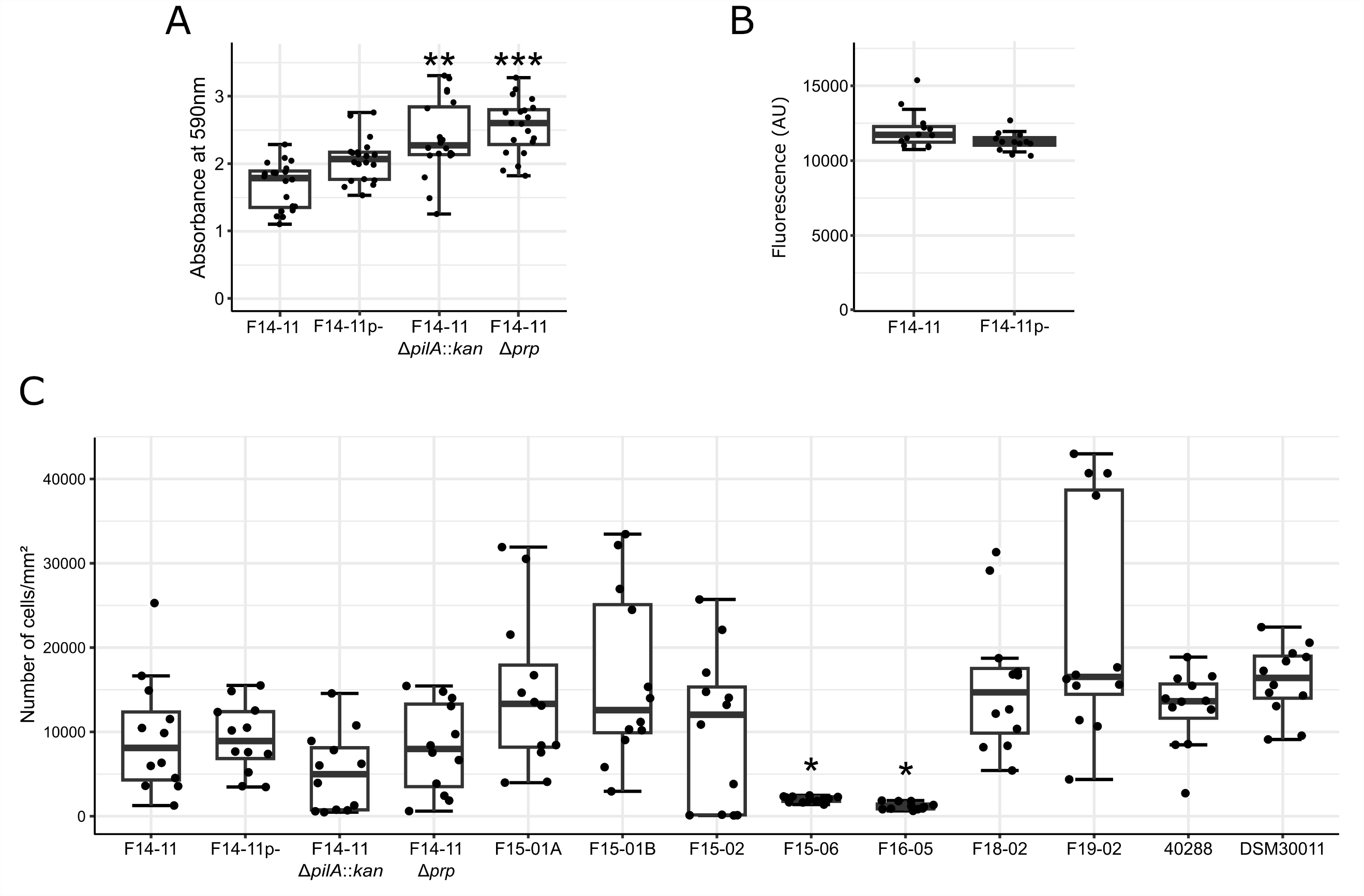

**FIG S5.** Adhesion and biofilm quantification. (A) Quantification of biofilm formation by F14-11 derivatives impaired for Type IV *pili* (∆*pilA*::*kan*) or Prp type I *pili* (∆*prp*) with crystal violet straining. (B) Congo red binding quantification of F14-11 wild-type and plasmid-less strains. (C) Adhesion quantification by the study strains, the dog UTI strain 40288 and the strong-adherent environmental strain DSM30011 after 1 hour of incubation using automatic cell counting in microscopy images. Error bars are generated from 12 replicas from 4 independent experiments. Statistically significant differences with F14-11 (Kruskal-Wallis rank-sum tests with Bonferroni-Holm correction) are highlighted with asterisks: ( * ) if the p-value < 0.05, ( * * ) if the p-value ≤ 0.01, and ( * * * ) if the p-value ≤ 0.001. The absence of asterisk indicates non-significant statistical difference.

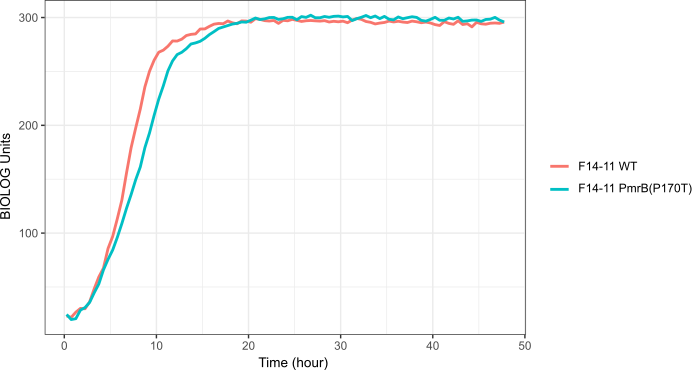

**FIG S6.** Growth curve of wild type F14-11 and its PmrB(P170T) derivative in rich medium (Biolog GenIII, control well).

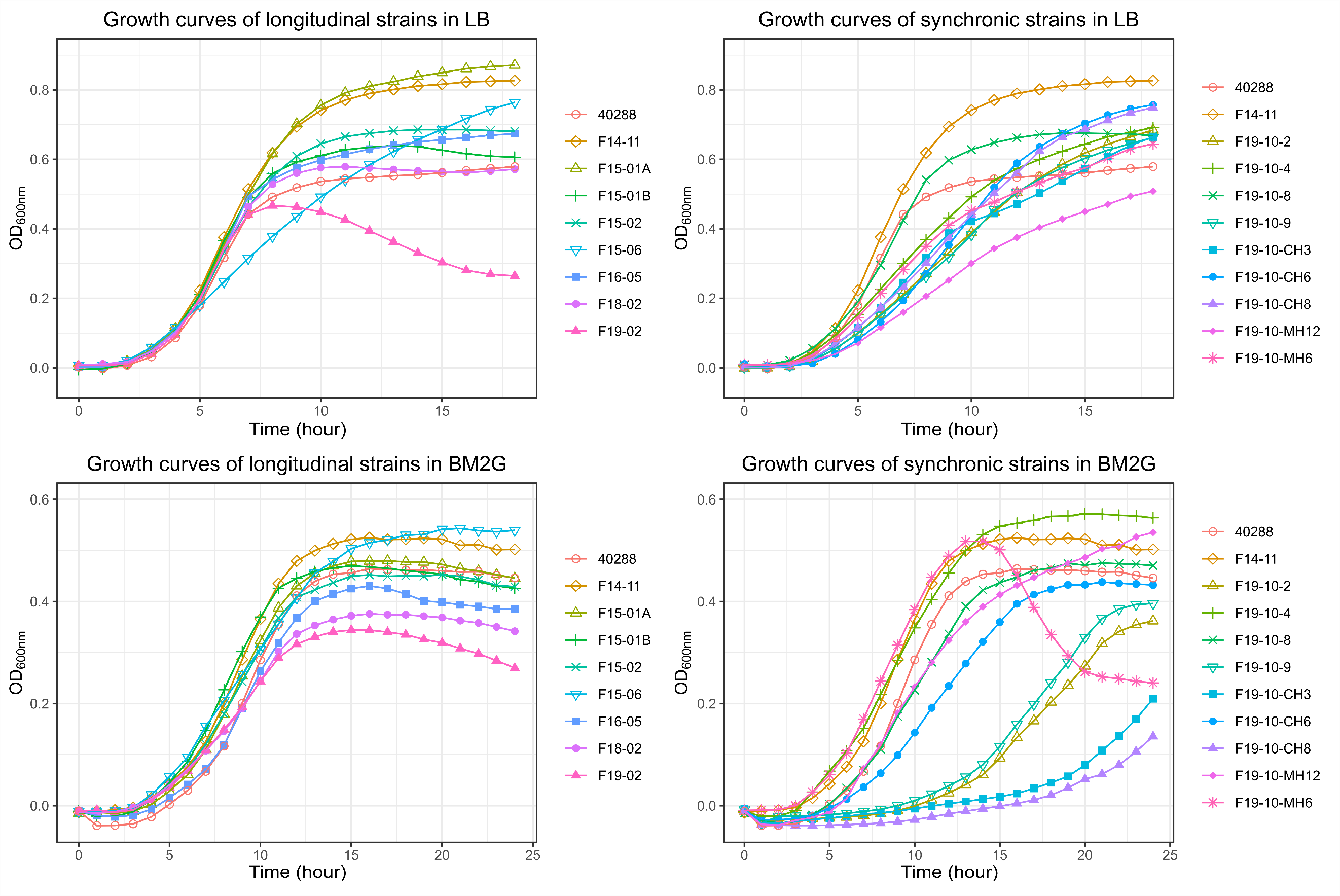

**FIG S7.** Growth rates of the longitudinal and synchronic strains in two different media (LB and BM2G).

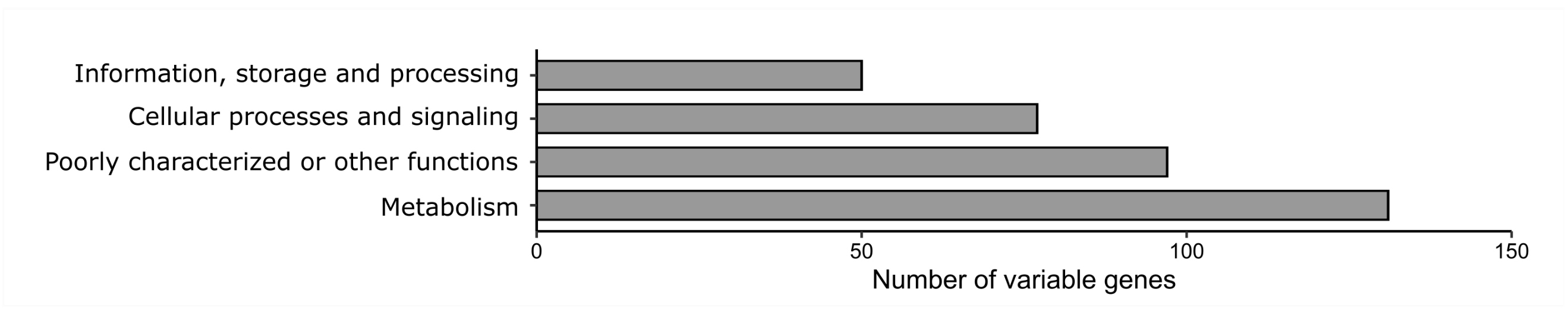

**FIG S8.** Distribution of functional categories of all the variable genes attributed by EggNOG-mapper v2 (5) in comparison to F14-11.
